# Draft genome of *Paraperlucidibaca* sp. CMO1 isolated from marine oil snow in the Churchill Marine Observatory, MB, Canada

**DOI:** 10.64898/2026.11.02.763556

**Authors:** Thamali Vidanage, R. Eric Collins

## Abstract

An oil-degrading enrichment culture was inoculated from Marine Oil Snow (MOS) collected during an oil-spill experiment at the Churchill Marine Observatory in winter 2024. The complete metagenome-assembled genome for *Paraperlucidibaca* sp. CMO1 was reconstructed, exhibiting the genetic potential to degrade hydrocarbons, including alkanes and aromatic compounds, as well as the ability to produce surfactants.

## 1 Genome announcement

*Paraperlucidibaca* is a genus of Gram-negative, rod-shaped bacteria belonging to the class Gammaproteobacteria (Oh et al., 2011). *Paraperlucidibaca* was first proposed in 2011 based on the notable differences in phenotypic properties with its closest member, *Perlucidibaca* (Oh et al., 2011). *Paraperlucidibaca* is a well-known hydrocarbon degrader in oil-contaminated seawater, exhibiting a significant enrichment in the presence of nutrients (Murphy et al., 2021). Here, we report the complete metagenome-assembled genome of *Paraperlucidibaca* sp. CMO1, isolated from the Ocean-Sea Ice Mesocosm (OSIM) facility at the Churchill Marine Observatory, Manitoba, Canada, during a winter 2024 oil-spill experiment.

A total of 114 L of Churchill Diesel oil was added to the experimental pool at the OSIM on December 3^rd^, 2024. Light brown flocs were observed floating on the pool’s surface 55 days after oil was added, which we refer to as Marine Oil Snow (MOS). The MOS particles were collected in aseptic glass jars and grown on Bushnell-Haas agar (plus 3% w/v NaCl) plates with motor oil spotted on the lid to use as the sole carbon source for bacterial growth (Malavenda et al., 2015). Cultures were maintained at 4 °C, and subcultures were obtained through streak-plating, but pure cultures were not obtained after multiple streaking, so analysis proceeded with the mixed culture. Genomic DNA extraction of the enrichment culture was performed using the DNeasy PowerSoil kit (Qiagen). Multiplexed DNA libraries were prepared using the Rapid PCR Barcoding Kit 24 v14 (SQK-RPB114.24) and loaded onto a FLO-MIN114 flowcell and sequenced on the Oxford Nanopore Technologies GridION device according to the manufacturer’s instructions (Oxford Nanopore Technologies, Oxford, UK). Twelve enrichment cultures were co-assembled using Flye 2.9.4-b1799 in metagenomics mode with a minimum overlap of 1000 bp. The co-assembly was polished in Flye using the built-in polisher, and a single circular contig containing the *Paraperlucidibaca* genome was reconstructed from the resulting assembly graph using Bandage v0.8.1 (Wick et al., 2015). Genome annotation was performed using RASTtk via the BV-BRC annotation service.

The complete genome is 2,526,375 bp in length, with a G+C content of 52.39% and 2,577 predicted protein-coding sequences, 45 tRNAs, and 4 rRNA genes (Table 2; Figure 1). Average Nucleotide Identity (ANI) of 98.37% with uncultured *Paraperlucidibaca* sp. (GCA_963974805.1) and 90.09% with strain *Paraperlucidibaca baekdonensis* DSM 26022.

**Figure 1.**
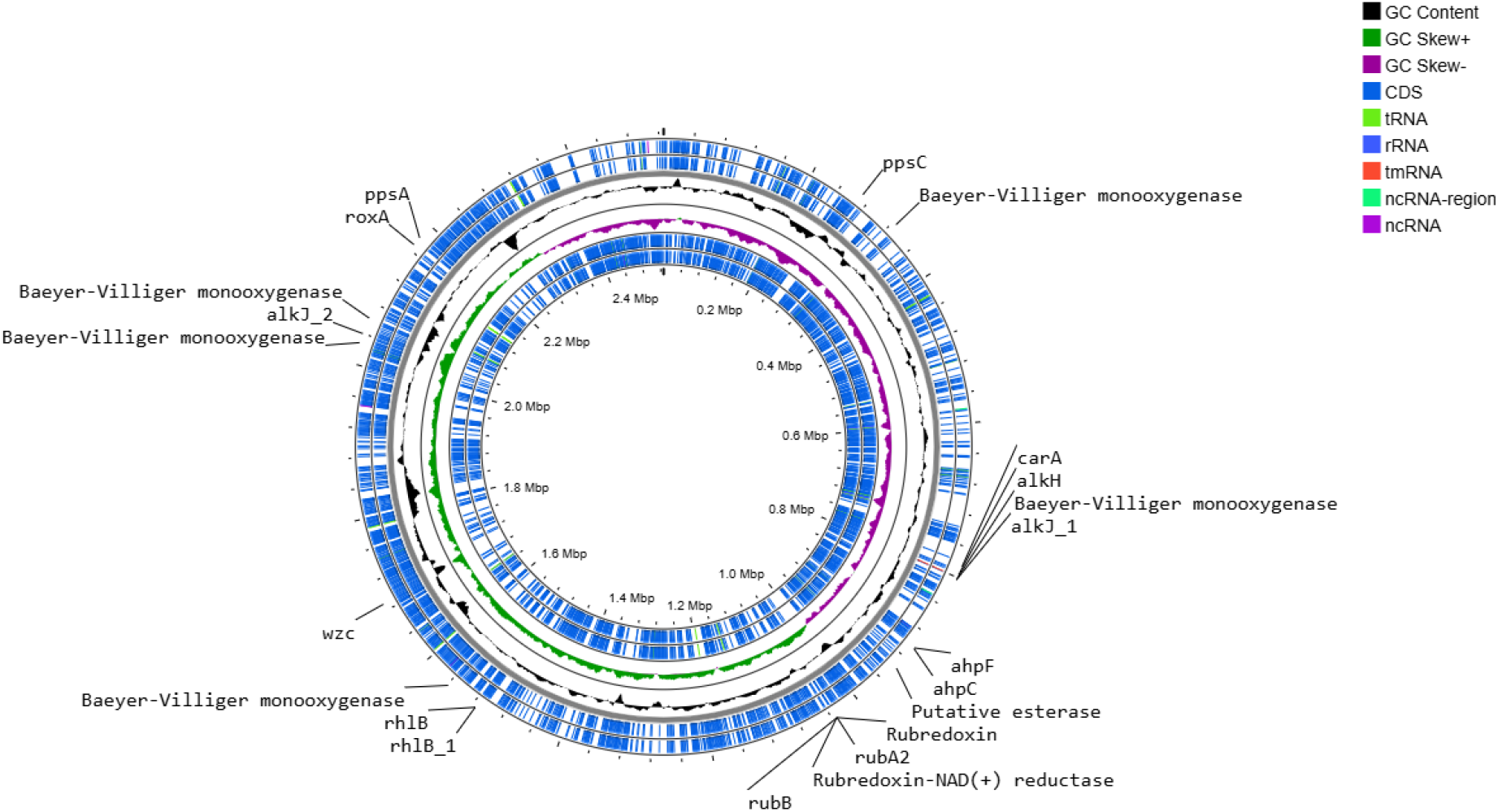
Genome circular diagram of *Paraperlucidibaca* sp. generated by Proksee (https://proksee.ca/).

Genes associated with hydrocarbon degradation were identified by querying the genome against the Hydrocarbon Aerobic Degradation Enzymes and Genes (HADEG) (Rojas-Vargas et al., 2023) and Calgary approach to ANnoTating HYDrocarbon degradation genes (CANT-HYD) (Khot et al., 2022) databases (Table 1). A noise cutoff was applied to CANT-HYD annotations to retain high-confidence gene hits. Functional gene annotation using HADEG was performed with thresholds set to --identity = 50 and --conn = 0.3, yielding a 94.68% true-positive rate and a 20.80% false-negative rate. Manual annotation using Prokka and Bakta revealed that the genome also encodes rubber oxygenase (*roxA*) (Cui et al., 2023), as well as genes associated with biosurfactant production, including rhamnolipids (*rhlB*) (Kumar et al., 2024), emulsan (*wzc*) (Nayarisseri and Singh, 2023), and plipastatin (*ppsC*) (Wang et al., 2024).

**Table 1.** Genes related to hydrocarbon biodegradation identified in the metagenome-assembled genome based on BV-BRC annotation and matches to curated hydrocarbon-degradation databases (HADEG and CANT-HYD). Genes retained following CANT-HYD noise cutoff filtering are indicated with an asterisk (*).

| Target name | Gene name | Description | CANT-HYD E-value | Availability in CANT-HYD | HADEG E-value | Availability in HADEG | BLAST hit E-value |
|---|---|---|---|---|---|---|---|
| fig\|2708021.11.peg.115 | *est* | Arylesterase precursor | - | - | 7.80e-57 | √ | W6R2Y2\|est<br>1.25e-75 |
| fig\|2708021.11.peg.360 | *AlmA_GroupI, AlmA_GroupIII* | Flavin-binding alkane monooxygenase | 3.3e-43 | √ | - | - | A5H9N6\|almA A5H9N8\|almA<br>1.29e-37 |
| fig\|2708021.11.peg.812 | *alkH_ald* | Aldehyde dehydrogenase | - | - | 8.27e-137 | √ | P12693\|alkH_ald<br>1.71e-175 |
| fig\|2708021.11.peg.816 | *AlmA_GroupI, AlmA_GroupIII* | Flavin-binding alkane monooxygenase | 6e-44 | √ | - | - | G1C7M4\|almA1.91e-36 |
| fig\|2708021.11.peg.915 | *ahpF* | Alkyl hydroperoxide reductase protein F | - | - | 2.68e-203 | √ | Q9I6Z2\|ahpF<br>0.0 |
| fig\|2708021.11.peg.916 | *ahpC* | Alkyl hydroperoxide reductase protein C | - | - | 2.77e-71 | √ | Q02UU0\|ahpC<br>9.78e-91 |
| fig\|2708021.11.peg.1041 | *alkG_rubA2_rdx* | Rubredoxin | - | - | 1.14e-26 | √ | Q9HTK8\|alkG_rubA2_rdx<br>3.1e-29 |
| fig\|2708021.11.peg.1042 | *alkT_rubB_rdxR* | Rubredoxin-NAD (+) reductase | - | - | 9.70e-120 | √ | Q0VTB0\|alkT_rubB_rdxR<br>3.15e-154 |
| fig\|2708021.11.peg.1585 | *BVMO* | Cyclohexanone monooxygenase | - | - | 5.07e-144 | √ | Q9I3H5\|BVMO<br>9e-180 |
| fig\|2708021.11.peg.1585 | *AlmA_GroupI, AlmA_GroupIII* | Flavin-binding alkane monooxygenase | 5.2e-45 | √ | - | - | K0CFH5\|almA E3UUD9\|almA<br>4e-36 |
| fig\|2708021.11.peg.2019 | *AlmA_GroupI, AlmA_GroupIII* | Flavin-binding alkane monooxygenase | 1.9e-238 | √ | 6.06e-171 | √ | A5H9N6\|almA A5H9N8\|almA A5H9P0\|almA E3UUD9\|almA G1C7M4\|almA GOM1\|almA K0CFH5\|almA Q6F7T9\|almA<br>0.0 |
| fig\|2708021.11.peg.2076 | *AlmA_GroupI, AlmA_GroupIII* | Flavin-binding alkane monooxygenase | 1.4e-40 | √ | - | - | A5H9N6\|almA<br>4.81e-35 |
| fig\|2708021.11.peg.2080 | *carAc* | Ferredoxin, 2Fe-2S | - | - | 2.62e-23 | √ | D5IGG4\|carAc<br>1.97e-30 |
| fig\|2708021.11.peg.2081 | *CYP153\** | Alkane-oxidizing cytochrome P450 | 8.5e-201 | √ | 1.68e-189 | √ | L7QFU8\|CYP153<br>0.0 |
| fig\|2708021.11.peg.2086 | *CYP153\** | Alkane-oxidizing cytochrome P450 | 3.8e-201 | √ | 1.75e-245 | √ | L7QFU8\|CYP153<br>0.0 |
| fig\|2708021.11.peg.2086 | *CYP450* | Putative cytochrome P450 hydroxylase | - | - | 1.75e-245 | √ | Q6RCE3\|CYP450 D0Q1H3\|CYP450<br>0.0 |
| fig\|2708021.11.peg.2109 | *AlmA_GroupI, AlmA_GroupIII* | Flavin-binding alkane monooxygenase | 1.2e-52 | √ | - | - | K0CFH5\|almA<br>1.87e-44 |
| fig\|2708021.11.peg.2119 | *carAc* | Ferredoxin, 2Fe-2S | - | - | 2.85e-24 | √ | D5IGG4\|carAc<br>3.06e-32 |
| fig\|2708021.11.peg.2120 | *CYP153\** | Alkane-oxidizing cytochrome P450 | 1.3e-201 | √ | 8.40e-190 | √ | L7QFU8\|CYP153 D0Q1H3\|CYP450 Q6RCE3\|CYP450<br>0.0 |
| fig\|2708021.11.peg.2168 | *AlmA_GroupI, AlmA_GroupIII* | Flavin-binding alkane monooxygenase | 2.3e-213 | √ | - | - | A5H9N6\|almA A5H9N8\|almA A5H9P0\|almA E3UUD9\|almA G1C7M4\|almA GOM1\|almA K0CFH5\|almA Q6F7T9\|almA<br>0.0 |
| fig\|2708021.11.peg.2316 | *AlkB\** | Alkane hydrolase | 4.9e-208 | √ | 6.17e-142 | √ | C0LMW6\|alkB<br>2.01e-141 |

**Table 2.** Annotation statistics of *Paraperlucidibaca* sp.

| **Features** | ***Paraperlucidibaca* genome** | **Community co-assembly** | **Original reads** |
|---|---|---|---|
| Genome length (bp) | 2526375 | 45525428 | 897915959 |
| No. of contigs/reads | 1 | 2864 | 479225 |
| No. of coding DNA sequences | 2577 | - | - |
| N50 (bp) | 2526375 | 33629 | 4344 |
| L50 (no. of contigs/reads) | 1 | 223 | 60354 |
| GC content (%) | 52.39 | 46.48 | 44.13 |
| tRNA | 45 | - | - |
| rRNA | 1 | - | - |
| Proteins with functional assignments | 1640 | - | - |
| Proteins with EC number assignments | 715 | - | - |
| Proteins with Pathway assignments | 530 | - | - |
| Proteins with GO assignments | 606 | - | - |
| Proteins with Subsystem assignments | 826 | - | - |

## Data availability

Sequence data from this study have been deposited at NCBI under BioProject PRJNA1176269 (https://www.ncbi.nlm.nih.gov/bioproject/PRJNA1176269).

## Author contributions

Thamali Vidanage – Conceptualization, Data curation, Formal analysis, Investigation, Methodology, Visualization, Writing – original draft, Writing – review & editing, Project administration R. Eric Collins – Conceptualization, Methodology, Writing – review & editing, Visualization, Supervision, Validation, Funding acquisition, Project administration

